# Sound source history warps perceived azimuth and elevation

**DOI:** 10.1101/2025.10.25.681944

**Authors:** Tanya Wen, W. Owen Brimijoin, Antje Ihlefeld

## Abstract

In real-world auditory localization, humans make decisions based on acoustic cues that reach the ear as well as from expectations of where the sound may originate. Past studies have shown that the auditory system makes rapid adjustments in response to changes in the statistics of recent stimulation, which help maintain sensitivity over the range where most stimuli occur. In the current study, we explored the auditory system’s adaptability and plasticity by presenting sounds in different spatial contexts and examined the subsequent effects on sound localization accuracy. Identical sounds were rendered at different azimuth and elevation ranges of [−15°, −30°, 0°] or [15°, 30°, 0°] presented in contextual blocks. Sounds presented at 0° azimuth exhibited a rightward shift in perception when the targets were blocked within the left hemispace, and a leftward shift when targets were blocked in the right hemispace. Sounds that were blocked in the bottom hemispace showed an upward bias in perceived elevation, although the reverse was not found to be significant. Furthermore, participants who showed more errors in overall localization showed a higher degree of bias. Our results corroborate the account that the mapping of auditory space is dynamic and shifts depending on the context in which sounds are presented.

## Introduction

Sound localization is a difficult task. The auditory system combines information from binaural cues, including interaural time differences (ITD) and interaural level differences (ILD) for sources in the horizontal plane (i.e., azimuth angle), as well as monaural spectral cues generated by the head-related transfer function (Middlebrooks & Green, 1991) for locations in the vertical plane (i.e., elevation angle). While these cues are reliable in anechoic conditions, everyday environments contain background sound sources and reverberation, resulting in ambiguity. Converging evidence suggests that instead of trying to compensate for these inherent ambiguities, people continuously update how they map audio cues to spatial location, warping the audio-spatial topology over time as they weigh new sensory and statistical evidence (Keating & King, 2015; Lingner et al., 2018).

As a result, the auditory system exhibits systematic errors, or biases, when localizing sounds. These biases can be either (1) static, that is, inherent to human perception, or (2) adaptive, such that it changes over time depending on stimulus history. Static biases have been linked to incomplete coordinate transformations within the auditory system, which is necessary to compensate for head rotation during audio localization (Freeman et al., 2017; Leung et al., 2008) or to facilitate interactions with the visual system (Lewald & Ehrenstein, 1998; Razavi et al., 2007; Willett et al., 2019). In the horizontal plane, pre-existing biases auditory localizations have been reported to be biased away from the center, such that listeners tend to overestimate the sound source to be at a greater angle away from their acoustic midline (Odegaard et al., 2015; Parise et al., 2012). These errors increase as the sound source is presented further into the periphery (Garcia et al., 2017; Makous & Middlebrooks, 1990), as characterized by the sigmoidal psychometric function linking source azimuth with perceived laterality. In a parallel observation, sounds rotating around the head appear to move faster at the front than at the side (Brimijoin, 2018), and these biases observed for motion estimation qualitatively match those biases observed for overestimation of static azimuths. Put together, these perceptual distortions of auditory space can be characterized as a relative expansion of space in the front and compression at the side. Biases in the vertical plane have been less studied, although some have suggested that auditory perceptions are found to be “compressed” or biased to the horizon (Carlile et al., 1997; Dobreva et al., 2011). For sound sources on the horizon (elevation = 0°), elevation perception tends to be biased upwards by a few degrees (Carlile et al., 1997; Parise et al., 2014) for most frequencies as well as for broadband sound. These biases increase when the sensory information in the peaks and notches of the HRTF becomes unreliable, such as the deterioration of sensitivity in higher frequencies in aging or presentation of a narrowband stimulus (Dobreva et al., 2011; Parise et al., 2014).

The auditory system is highly adaptive (King et al., 2001; King et al., 2011). In order to adjust to a rapidly changing world, the brain must continuously update its neural representations to reflect changes in sensory input and behavioral goals (Keating & King, 2015). Plasticity in auditory space calibration has been shown in a wide range of conditions (Mendonça et al., 2014), including continuous passive exposure to altered sounds (van Wanrooij & van Opstal, 2005; Carlile & Blackman, 2014) and as well as from training (Shinn-Cunningham et al., 1998; Majdak et al., 2013). Studies showed that participants can relearn the mapping of auditory cues to spatial locations after alterations to the shape of the pinnae (Hoffman et al., 1998) and unilateral ear plugging (Kumpik et al., 2010; Zonooz & Van Opstal, 2019). Common across these studies, after restoring to their original hearing condition, participants immediately regain their pre-altered localization performance (Irving & Moore, 2011). This reflects that auditory space mapping is flexible and adaptive, such that different patterns of spectral or binaural features can map to the same location in space (Grothe et al., 2010; Stitt et al., 2019).

Adaptive biases can occur when the short-term statistics of the preceding stimulus distribution biases localization performance (Carandini & Heeger, 2011; Getzmann, 2004; Maier et al., 2012). This is established through a process called adaptive coding, where neurons adjust their tuning properties to better represent the prevailing stimulus environment (Dean et al., 2005; Hildebrandt et al., 2015; Stange et al., 2013). For example, by presenting sounds in spatial ranges from narrow to wide, participants were able to increase precision by rapidly changing their response sensitivity to the experimental distribution of the targets (Ege et al., 2019). Psychophysical assessments show that the auditory system adapts to the mean of the ITD and ILD distribution, leading to a misjudgement of sound source laterality, such that stimuli tend to be perceived in a position shifted in the opposite direction of the distribution they were adapted to. On the horizontal plane, stimuli were more likely to be perceived as located to the left of the midline when the mean stimulus distribution is shifted to the right, and vice versa (Alamatsaz & Ihlefeld, 2019; Dahmen et al., 2010; Kashinob & Nishida, 1998; Kopco et al., 2007). These studies suggest that the warping of the audio-spatial map is not constant, but dynamically changes over time as the listener continuously recalibrates to the statistics of their environment.

While it has been demonstrated that auditory source localization is susceptible to both static and adaptive biases, it is incompletely understood how the mapping of spatial audio cues to locations changes over time. In the current study, we varied the short-term stimulus history by presenting sounds in blocks of locations restricted to a certain hemispace. We predicted that this contextual blocking would skew the perception of auditory sources (Alamatsaz & Ihlefeld, 2019; Dahmen et al., 2010) in both azimuth and elevation. However, we cannot directly compare azimuth and elevation, as the latter is additionally impacted by asymmetrical floor reflections. The second goal of the study is to explore whether the amount of systematic error that is observed during contextual blocking is related to individuals’ localization ability. Studies have suggested that listeners change their reliance on auditory signals and prior biases as a function of their relative reliabilities or signal-to-noise ratio. In particular, the reliance on priors increases with stimulus ambiguity (Garcia et al., 2017; Körding & Wolpert, 2006). We predict that localization biases from contextual blocking may be increased in poorer listeners. Finally, we extend on prior work by using virtual reality (VR). Prior work using loudspeakers is limited by the inability to disentangle reverberation in the test environment from the acoustics reaching the listener’s ears. Alternatively, headphone experiments are usually conducted with headlocked audio. By using VR, we are able to present world-locked anechoic sounds to test localization performance.

## Methods

### Participants

Thirty-two participants were recruited from the community to participate in this study at Meta Reality Labs in Redmond, WA. All participants were naive to the study design. While we targeted recruitment for participants under the age of 45 to minimize the effect of age, we had a small number of older participants show up to the study. All participants were neurologically healthy, with normal or correct-to-normal vision, and have self-reported normal or close-to-normal hearing. These participants also had their Head-Related Transfer Function (HRTF) previously measured at our facilities. Procedures were carried out in accordance with ethical approval obtained from the Institutional Review Board, and participants provided written, informed consent before the start of the experiment. Participants were compensated for their time. Three participants were excluded due to technical errors or opting out of the study, which resulted in incomplete data. Another five participants were excluded due to poor performance or lack of spatial sensitivity. The exclusion criteria involved not perceiving the most extreme azimuth or elevation (i.e., −/+ 30°) within their respective hemispace in at least one or more blocks.

The percentage of non-sensitive listeners was similar to our previous experiences. Additional analyses verified that inclusion of non-sensitive listeners did not change the overall results. This resulted in a total of 24 participants (12 male, 12 female; ages 25-60, mean = 35.79, SD = 8.38; 23 right-handed, 1 handedness not provided). The data and code have been made publicly available and can be accessed at https://github.com/facebookresearch/AuditoryPerceptionRangeBias.

### Task Procedure

At the beginning of the experiment, participants underwent a short practice session to familiarize participants with spatial audio as well as using the virtual reality (VR) headset and controllers. During the practice, participants were aligned to the center of the sphere at the beginning of each trial. On each trial, they listened to a 3-second click train noise and were instructed to localize the position of the sound. Participants were asked to face forward, but were allowed to make small head movements (up to ∼10° yaw/pitch/row) while the sound was playing, as if they were naturally listening and attending to the sound, but not big movements. After they have located the sound, participants were asked to reach out with their closest hand as if they were grabbing the sound. Once they had located the position of the sound, they were asked to press the “X” or “A” button on the left or right controllers to indicate their response. After the button was pressed, a green dot would appear at the sound’s ground truth location. To encourage participants to learn the association of the sound and its location, participants were asked to look at the green dot and re-orient their controllers to the green dot’s location and make another button press on the controllers to confirm the location. Participants were further instructed to use the green dot as feedback, and use that feedback to improve future performance.

The instructions for the main experiment were similar to that of the practice. Here, on each trial, participants aligned themselves to the center of the sphere, and listened to a 2-second pink noise sound while only making small movements with their head. Afterwards, participants pointed the controller at the sound location and recorded their response. Here, participants were not given feedback on their performance, but were asked to perform as accurately as possible. The main experiment was presented in five different contextual blocks, where we tested (1) the full range of manipulated azimuths and elevations, (2) azimuths blocked such that all stimuli were presented in the (a) left hemispace and (b) right hemispace, (3) elevations blocked such that all stimuli were presented in the (a) bottom and (b) top hemispace. The full range context was presented first and served as a baseline. The order of the remaining four contexts were counterbalanced across participants. Participants were given mandatory breaks of at least 5 minutes between the practice and between each block to minimize carryover effects of the contextual adaptations. Participants were not explicitly informed of the blocking manipulation.

### Materials and Stimuli

Sounds were presented at 75 dB SPL through DT990 headphones, ensuring audibility without causing discomfort. Prior to testing, the DT990 headphones were equalized using the Brüel & Kjær HATS 5128-B mannikin, Brüel & Kjær 1704-A-002, and RME Fireface UFX II sound card; and the inverse filter resulting from the equalization was applied to the audio. The audio was rendered through the headphones connected to the Quest 3 headset. Spatial audio was generated by convolving each participant’s individual HRTF with the audio file using an internal convolution and spatialization tool. All sounds were rendered at 2 meters distance from the head.

For practice, we generated 30 noise tokens between 200 Hz to 16,000 Hz with a duration of 3 seconds with 60 ms on/off ramps. These were a train of clicks at 100 Hz with normally distributed random phase, similar to that described in Brungart and Simpson (2008). The output was roved in 1/3 octave bands with 3 bands per octave. A gain is applied with uniform distribution within +/-3 dB for each band. The practice sounds were randomly drawn on each trial distributed around the entire sphere. The locations were jittered [−5° to 5°] around azimuths = [45°, 135°, 225°, 315°] and elevations = [−70°, −40°, −10°, 10°, 40°, 70°]. This generated 48 training trials. We additionally added 12 trials of random locations drawn from azimuths = [0° to 359°] and elevations = [−80° to 80°]. We did not go further beyond +/-80° elevation due to the difficulty of reaching directly under the chair to point at extreme elevations. There were 60 training trials in total.

In the main experiment, the target sound consisted of pink noise, and had a duration of 2 seconds, with 10 ms on/off ramps. The cut-off frequencies of the noise equaled 100 to 18,000 Hz. We generated 100 pink noise tokens and randomly presented one on each trial. We rendered sounds at locations of combinations azimuth = [−30°, −15°, 0°, 15°, 30°] and elevations = [−30°, −15°, 0°, 15°, 30°] for the full range test. Each unique position appeared twice to the participants in random order, resulting in a total of 50 trials. We decided to have a smaller range of elevation as piloting suggested that lower elevations, especially near the center, were more physically difficult to reach, due to the position of the chair and the participants’ legs when seated. For the azimuth blocking, we presented stimuli at [−30°, −15°, 0°] (left hemispace) and [0°, 15°, 30°] (right hemispace) at all [−30°, −15°, 0°, 15°, 30°] elevations. For elevation blocking, stimuli were presented at [−30°, −15°, 0°] (bottom hemispace) and [0°, 15°, 30°] (top hemispace) at each of the [−30°, −15°, 0°, 15°, 30°] azimuths. Within each context, each unique condition was presented a total three times for each participant, resulting in a total of 45 trials per block. Each block took participants approximately 6-7 minutes to complete.

Participants were seated in a sound-attenuated quiet room during the study wearing the Quest 3 headset. The task was rendered in virtual reality (VR) through Unity. Inside the VR environment, participants were situated inside a gray sphere with a 1.5 meter radius, which was calibrated at the start of the experiment such that it was centered at the participant’s head. An alignment GUI was visible to the participants, showing real-time feedback of their head position from the sagittal and transverse viewpoints. Participants were instructed to get into alignment before the start of each trial. We recorded thumb-pointing responses with the Quest Pro controllers (which have inside-out camera-based tracking, and were separately validated to accurately track location). Participants were instructed to fully reach out with the controllers towards the source of the sound, to indicate their perceived location. Furthermore, they were instructed to point with the controller that is closest to the target sound, and were allowed to use either hand if they perceived it as equally distant, such as when it is in their midline.

### Data analysis

The thumb pointing results were converted into spherical coordinates and analyses focused on perceived azimuth and elevation (Carlile, 1996; Poirier-Quinot et al 2022). A point directly in the front of the participant is described as 0° azimuth and 0° elevation. Negative azimuth values correspond to the left hemifield and positive azimuth values correspond to the right hemifield. Similarly negative elevation is below the horizon and positive is above the horizon. The current study did not focus on front-back confusions, so for the following analyses azimuths localized behind the participant were folded into the front (e.g., 150° would be analyzed as 30°). All analyses were performed in Matlab 2023b (Mathworks, Inc., Natick, MA, USA).

A linear mixed effects model (LME) was used to model the perceived azimuths and elevations. Although Alamtsaz & Ihlefeld (2019) used a non-linear mixed effects model with a sigmoid link function, our ranges fall within the area where perceived locations are mostly linear with the sources, and an LME would be most suitable. The model included fixed effects source azimuth (*α*_*y1*_), source elevation (*α*_*y2*_), and context (*α*_*y3*_), with context coded using a dummy variable. Random effects of individual differences and within-context trial number was used to model the intercept (β_y0_), which could be interpreted as the perceived midline. The equation is described as follows:

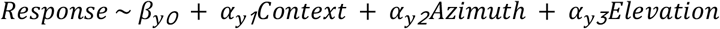

Next, we examined whether localization biases differed across different contexts. Biases were calculated by subtracting the perceived azimuth (or elevation) in the blocked context with the perceived azimuth (or elevation) in the full range context, and a positive bias indicates that the sound is perceived to the right (or above) the actual source. To examine the effect of azimuth blocking on localization biases, we performed two Context (full, left/right) × Azimuth (0°, −/+15°, −/+30°) repeated measures ANOVAs. The left and right contextual blocking were separately compared with the full context baseline condition separately. Similarly, two Context (full, bottom/top) × Elevation (0°, −/+15°, −/+30°) ANOVAs were performed to analyze the effect of blocking by elevation on biases. As 0° azimuth locations overlapped between the left and right contexts, and 0° elevation locations overlapped between the bottom and top contexts, we additionally focused on the 0° condition. The Greenhouse-Geisser correction was used to adjust for sphericity.

As we asked participants to point with their hand that is closest to the sound, and that either hand can be used when the sound is in the midline, it would be interesting to see whether their response hand differed depending on context. A one-way repeated measures ANOVA was used to examine the effect of azimuth blocking (full, left, right) and elevation blocking (full, bottom, top) on the proportion of right-hand responses at 0° azimuth and 0° elevation, respectively.

Finally, it is possible that participants who were less sensitive to spatial audio relied more on contextual range during the location judgements, therefore, a correlation was performed between performance and bias. Azimuth localization performance was calculated using the mean absolute error (MAE) of all source positions across the five contexts, leaving out 0° azimuth locations. Azimuth bias was calculated as the difference in bias between the left and right blocking contexts. The same method was used to calculate elevation performance, but leaving out 0° elevation locations. And elevation bias was calculated as the difference in bias between the bottom and top blocking contexts.

## Results

### Overall Accuracy

We calculated the average angle error across the five blocks (mean = 21.95°, SD = 11.68°) and found no significant correlation with participant age (Pearson’s r = 0.33, p = 0.12) or differences between genders (t = −0.96, p = 0.36, Cohen’s d = −0.41).

### Linear Mixed Effects Model

The perceived locations as a function of source azimuths and elevations are shown in Figure 1 A and C. In the full range context as well as the left and right blocking, when the source was in the periphery, azimuth perception showed a lateral bias, such that azimuths were overestimated. On the other hand, when the source was in the midline, azimuth localization was biased towards the contralateral hemispace in the left and right contextual blocks. Similar results were observed for elevation, where an overestimation of elevation was observed in the full, bottom, and top contexts; however, the horizon was overestimated when the context was blocked in the bottom compared to the top.

**Figure 1.**
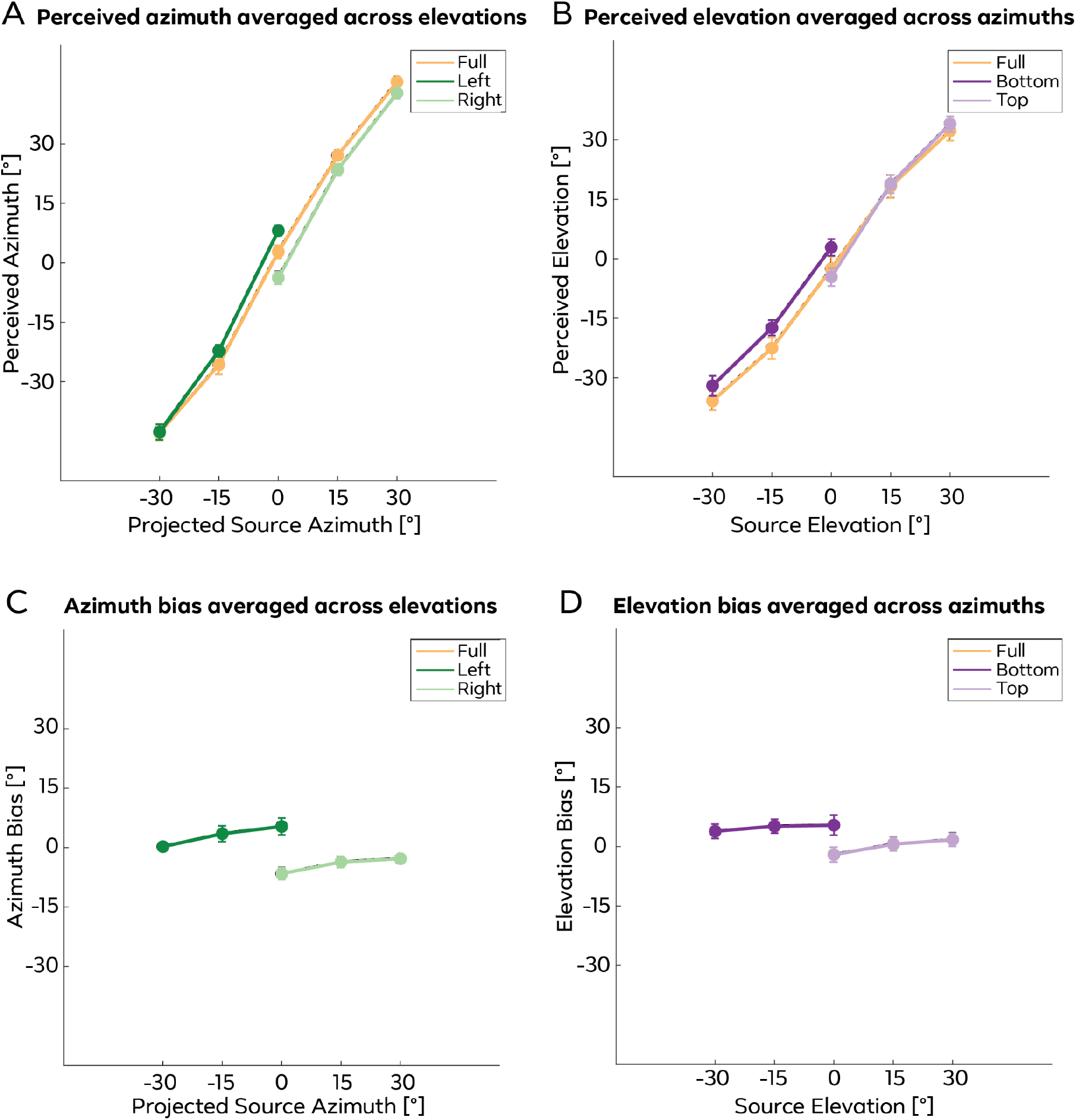
A. Perceived azimuth in the full (orange), left (dark green), and right (light green) contexts, as a function of source azimuth. B. Perceived elevation in the full (orange), bottom (dark purple), and top (light purple) contexts as a function of source elevation. C. Azimuth bias, defined as the difference between the left or right context and the full context, averaged across elevations plotted across each tested azimuth. The left context shows a negative bias at 0° azimuth, but positive biases at all other azimuths; on the other hand, the right context shows a positive bias at 0° azimuth, but negative biases at all other azimuths. D. Elevation bias, defined as the difference between the top or bottom context and the full context, averaged across azimuths plotted against each tested elevation. The bottom condition was more positively biased compared to the full condition. At 0° elevation, the bottom context showed a positive bias, while the top context showed a negative bias. The biases at 0° elevation were significantly different between the bottom and top contexts. Error bars represent standard errors.

As shown in Table 1, perceived azimuth significantly increased with source azimuth, as expected. Furthermore, the effect of target range in the left and right hemispaces were significant compared to the full condition baseline. On average, participants were biased 3.30° to the right when the targets are blocked within the left hemispace, and 6.30° to the left when the targets are blocked within the right hemispace.

**Table 1.** Effect of target range on perceived azimuth. Results of LME (24 participants), with fixed effects of target range, source azimuth, and source elevation; random effects of participant and trial number.

|  | Estimate | SE | tStat | pValue |
| --- | --- | --- | --- | --- |
| Intercept | 2.50 | 1.11 | 2.25 | 0.02 |
| azimuth | 1.67 | 0.02 | 71.69 | 0.00 |
| elevation | 0.00 | 0.02 | -0.11 | 0.91 |
| Range_Left | 3.30 | 0.97 | 3.39 | 0.00 |
| Range_Right | -6.30 | 0.97 | -6.47 | 0.00 |

Table 2 depicts the results of modeling perceived elevation on target range, azimuth, and elevation. As expected, perceived elevation significantly increased with source elevation. Furthermore, when the target range was in the bottom hemispace, elevation responses were biased 4.51° upward compared to the full range baseline. There was no significant effect on elevation when targets were presented in the top hemispace.

**Table 2.** Effect of target range on perceived elevation. Results of LME (24 participants), with fixed effects of target range, source azimuth, and source elevation; random effects of participant and trial number.

|  | Estimate | SE | tStat | pValue |
| --- | --- | --- | --- | --- |
| Intercept | -2.09 | 2.08 | -1.00 | 0.32 |
| azimuth | -0.03 | 0.01 | -1.83 | 0.07 |
| elevation | 1.20 | 0.02 | 61.11 | 0.00 |
| Range_Bottom | 4.51 | 0.82 | 5.51 | 0.00 |
| Range_Top | 0.22 | 0.82 | 0.26 | 0.79 |

### Effect of Context on Azimuth and Elevation

Figure 1C shows the degrees of azimuth bias, defined as the difference in the left and right contexts versus the full context, plotted against each tested azimuth. To compare the different contexts with the baseline condition on azimuth perception, we first conducted a Context (left vs. full) × Azimuth (0°, −15°, −30°) repeated measures ANOVA. Results showed a main effect of Context (F(1,23) = 4.59, p = 0.04, 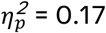) and Azimuth (F(2,46) = 345.45, p < 0.001, 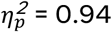). There was no significant Context × Azimuth interaction (F(2,46) = 2.35, p = 0.11, 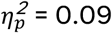). The main effect of Context was driven by sources in the left context being perceived overall more rightward compared to the full context (t = 2.14, p = 0.04, Cohen’s d = 0.44). Next, a Context (right vs. full) × Azimuth (0°, 15°, 30°) repeated measures ANOVA showed a main effect of Context (F(1,23) = 11.66, p < 0.01, 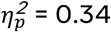) and Azimuth (F(2,46) = 429.67, p < 0.001, 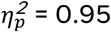). There was also a significant Context × Azimuth interaction (F(2,46) = 3.88, p = 0.04, 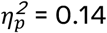). The main effect of Context was driven by the right context being perceived overall leftward compared to the full context (t = −3.41, p < 0.01, Cohen’s d = −0.70). The magnitude of the bias was significantly stronger at 30° than at 0° (t = −2.56, p = 0.02, Cohen’s d = −0.52).

A one-way repeated measures ANOVA comparing the perceived azimuths at 0° for the full, left, and right contexts showed a significant main effect of Context (F(2,46) = 15.83, p < 0.001,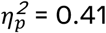). Post hoc paired t-tests suggests the three contexts were all significantly different from each other (left vs. full: t = 2.21, p = 0.04, Cohen’s d = 0.45; right vs. full: t = −3.84, p < 0.001, Cohen’s d = 0.78; left vs. right: t = 5.47, p < 0.001, Cohen’s d = 1.12). Put together, these results suggest that contextual blocking of azimuths affect participants’ perception of sound location. Furthermore, participants show a bias towards the opposite hemispace when the stimulus is presented at the midline, but a bias away from the midline when presented laterally.

Figure 1D shows the degrees of elevation bias, defined as the difference in the bottom and top contexts and the full context, plotted against each tested elevation. We performed the same analysis to examine the effect of context and elevation on elevation bias. A Context (bottom vs. full) × Elevation (0°, −15°, −30°) repeated measures ANOVA showed a significant main effect of Context (F(1,23) = 8.26, p < 0.01, 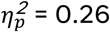) and Elevation (F(2,46) = 149.33, p < 0.001, 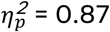). There was no Context × Elevation interaction (F(2,46) = 0.18, p = 0.82, 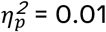). Post hoc analysis showed that the bottom context showed an overall more positive (i.e., upward) elevation bias (t = 2.87, p < 0.01, Cohen’s d = 0.59). The Context (top vs. full) × Elevation (0°, 15°, 30°) repeated measures ANOVA showed a main effect of Elevation (F(2,46) = 171.48, p < 0.001, 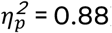), but no significant main effect of Context (F(1,23) < 0.01, p = 0.95, 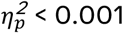) or Context × Elevation interaction (F(2,46) = 1.28, p = 0.28, 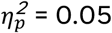).

A one-way repeated measures ANOVA comparing the elevation biases at 0° for the full, bottom, and top contexts showed a significant main effect of Context (F(2,46) = 5.05, p = 0.01, 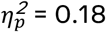). Post hoc t-tests showed a significant difference in bias between the bottom and top contexts at 0° elevation (t = 3.31, p < 0.01, Cohen’s d = 0.67), with the bottom context showing more positive (i.e., upward) bias compared to the top context. There were no significant differences between the bottom or top contexts with the full context (bottom vs. full: t = 1.91, p = 0.07, Cohen’s d = 0.39; top vs. full: t = −0.96, p = 0.34, Cohen’s d = −0.20). These results suggest that contextual blocking of elevation affects participants’ perception of sound location.

### Effect of Context on Azimuth and Elevation

The proportion of right-hand responses towards 0° azimuth and 0° elevation as a function of context is shown in Figure 2. A one-way repeated measures ANOVA with showed a significant main effect of Context when comparing the two azimuth blockings and the full context (F(2,46) = 18.43, p < 0.001, 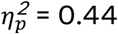). Post hoc comparisons of the three contexts showed that the proportion of right-hand responses were all significantly different across contexts. Specifically, participants were more likely to respond with their right hand towards 0° azimuth when sounds were predominantly presented in the left hemispace, compared to the full context (t = 3.28, p < 0.01, Cohen’s d = 0.67). On the other hand, participants were less likely to respond with their right hand when sounds were blocked in the right hemispace (t = −3.13, p < 0.01, Cohen’s d = −0.64). The same analysis was performed to examine the effect of elevation blocking on response hand. Results showed no significant main effect (F(2,46) = 0.29, p = 0.73, 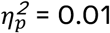).

**Figure 2.**
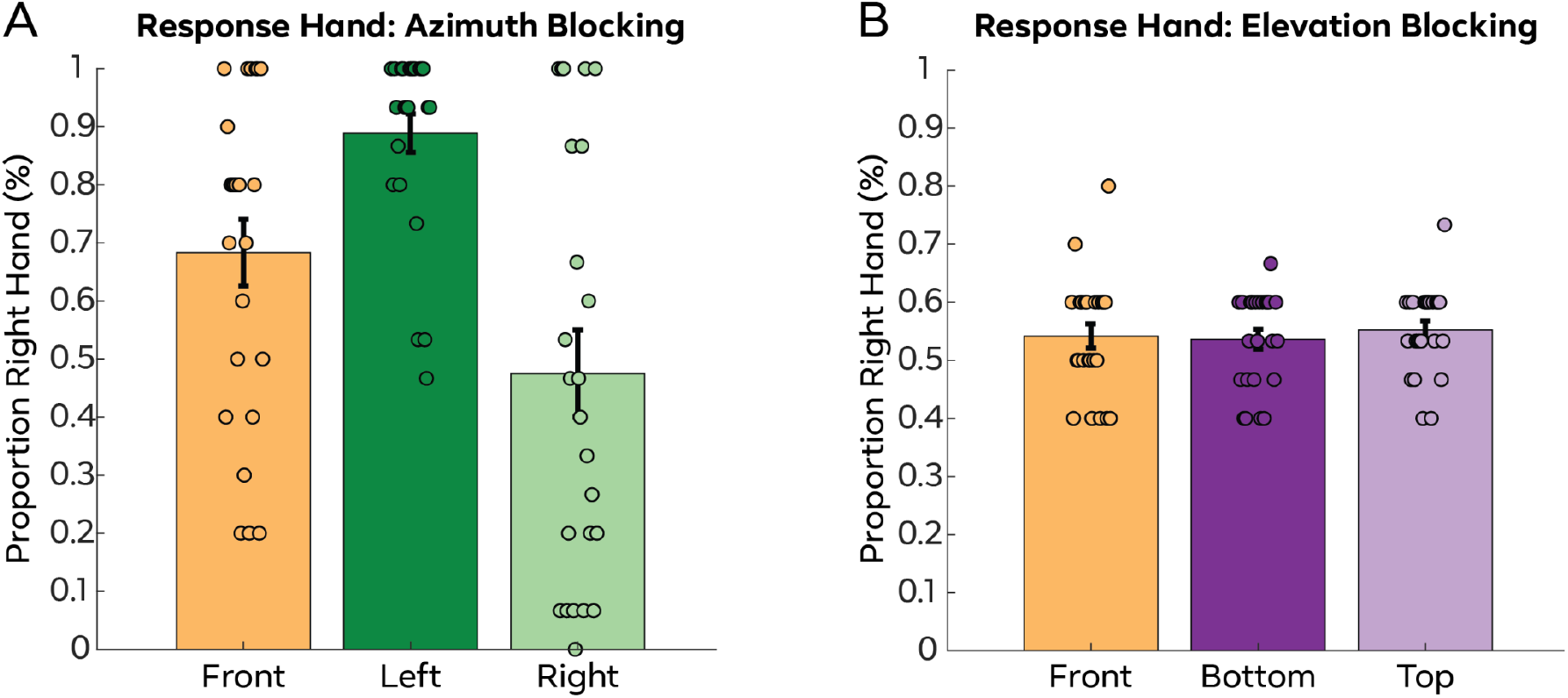
A. The proportion of right hand responses towards 0° azimuth in the full context, and when the sounds are predominantly blocked in the left or right hemispace. Participants are more likely to use their right hand in the left context, and less likely to use their right hand in the right context, compared to the full context. B. The proportion of right hand responses towards 0° elevation in the full context, and when the sounds are predominantly blocked in the bottom or top hemispace. No difference in response hand was found across the three conditions. Each dot represents each individual participant, with each individual staggered from left to right. The error bars represent standard error of the mean.

### Relationship Between Performance and Bias

Figure 3A shows the correlation between azimuth performance, measured by MAE, with the differences in azimuth 0° perception in the left versus right contextual block. Results showed a positive relationship (Pearson’s r = 0.56, p < 0.01), suggesting that participants with larger azimuth localization errors showed a stronger bias. Figure 3B depicts the correlation between elevation MAE with bottom versus top contextual differences at elevation 0°. The relationship was in the positive direction, however, did not reach statistical significance (Pearson’s r = 0.32, p = 0.13).

**Figure 3.**
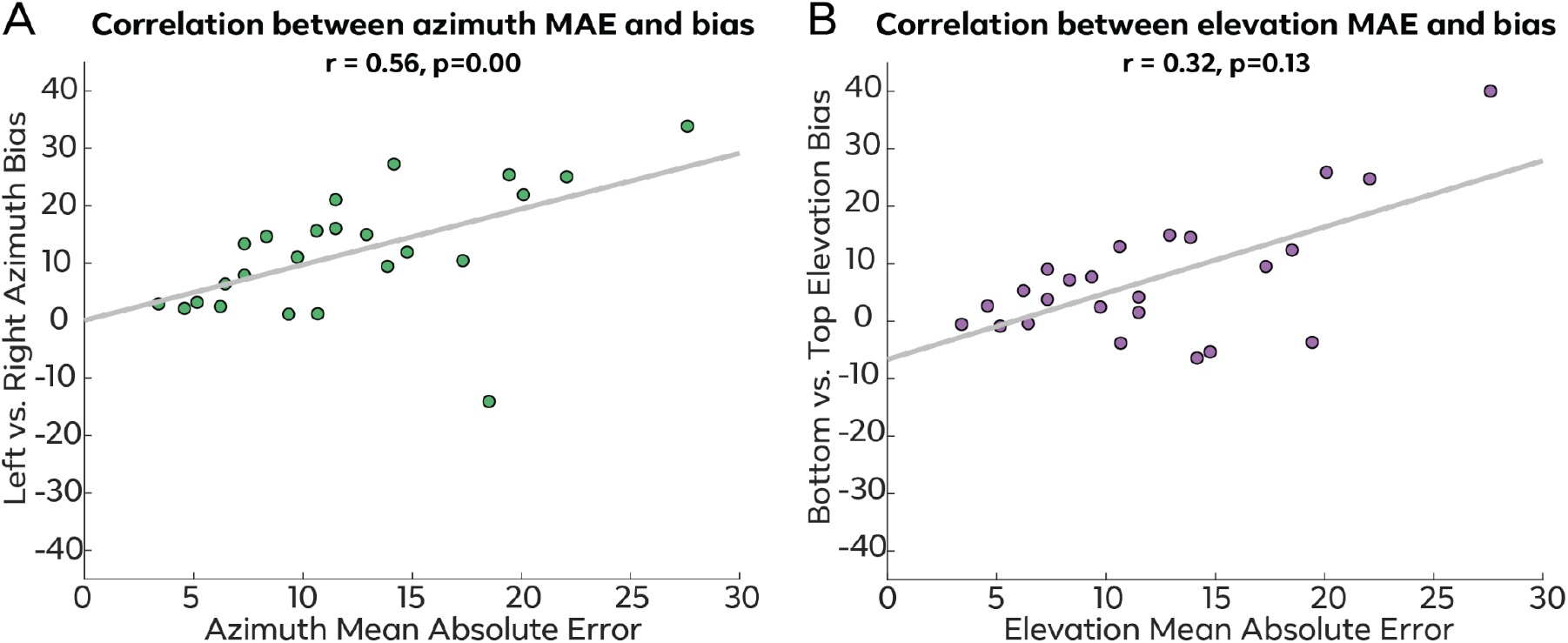
A. A positive relationship was found between the mean absolute error in azimuth localization and azimuth bias. B. Relationship between the mean absolute error in elevation localization and elevation bias did not reach statistical significance.

## Discussion

Our study showed that the spatial context in which targets are presented affect their perceived location. By presenting sounds in blocks predominantly in the left or right hemispace resulted in 0° azimuth locations showing a robust response bias towards the contralateral side. Furthermore, participants were more likely to use the contralateral hand to respond, compared to the full context. On the other hand, sounds that were presented laterally within those blocks (i.e., −/+ 15° and −/+30°) showed a response bias away from the center, such that listeners tend to overestimate the target location to be at a greater angle away from their acoustic midline. When examining the effect of blocking on elevation, we found that at 0° elevation locations, sounds presented in the bottom context showed an increased upward bias compared to that of the top context. Additionally, sounds presented in the bottom context showed an overall upward bias compared to the full condition. Finally, we showed that participants with larger azimuth errors in localization were more likely to be biased by target range.

Some previous studies have also reported perceptual biases during auditory localization, such that participants are biased towards the periphery when localizing auditory stimuli on the horizontal plane (Dobreva et al., 2011; Lewald & Ehrenstein, 1998; Odegaard et al., 2015). We observed overestimation of azimuths and elevations for eccentric sources in all contextual conditions. This systematic error in perception has been explained by some studies as an expansion of frontal space and a contraction at the sides (Brimijoin, 2018), or Bayesian prior for the periphery (Freeman et al., 2017; Odegaard et al., 2015).

Consistent with Bayesian decision theory, when sensory information is less reliable, participants would rely more on the prior when making localization judgements (Garcia et al., 2017; Körding & Wolpert, 2004, 2006). This is in line with our observation that participants who were less able to distinguish the target location exhibited greater biases. While this suggests that priors and uncertainty influence auditory localization biases, as priors biasing estimates to the periphery would become less strong as eccentricity increases (Garcia et al., 2017), we did not observe such a pattern (Figure 1A and B). Furthermore, one might expect priors to be updated by sensory evidence, such that they should converge towards the mean of the stimulus set within each blocking context over time. In this case, we would expect to see a bias towards 15° azimuth or elevation, but our results showed that participants instead expanded their perception in both directions outside the tested range, including showing a contralateral bias when sounds were presented at midline (Alamatsaz & Ihlefeld, 2019). Thus, the full pattern of our results cannot be explained by Bayesian priors alone.

We found that perceptual biases changed depending on how the sounds were blocked, as evident in how 0° was differently perceived in different contexts. The effect of context-dependency or stimulus range on perception has been studied in other contexts, including sensory judgements (Cheadle et al., 2014; Murai et al., 2016), reward (Bavard et al., 2021; Nieuwenhuis et al., 2005; Palminteri et al., 2015), and task difficulty (Wen & Egner, 2023). In these cases, the context provides a reference point to which stimuli can be compared before making a decision. For example, in economic valuation, 0¢ is high compared to −40¢ (lose money), but low compared to 60¢ (gain money; Nieuwenhuis et al., 2005). In our experiment, 0° azimuth would be more clockwise than −15° and −30° when stimuli are presented in the left hemispace, and more counterclockwise than 15° and 30° when targets are in the right hemispace. We suggest the relative spatial location within a context may contribute to the biases observed at midline. On a neuronal level, adaptation is a widespread property in which auditory neurons reduce their response towards the prevalent stimulus distribution, thereby enabling their responses to adapt and align with the constantly changing statistics of sounds that reach the ears (Dean et al., 2008; Willmore & King, 2023). Instead of maintaining an accurate representation of absolute sound-source locations, adaptation allows the auditory system to emphasize the relative differences between these locations, which can be a mechanism to increase discriminability between signals (Dahmen et al., 2010; Getzmann, 2004; Gleiss et al., 2019; Stange et al., 2013). As a consequence, such adaptation can cause a shift between the mapping of neuronal responses and sensory input, such that the perceived localization of sounds is shifted away from the adapting stimuli (Carlile et al., 2001; Lingner et al., 2018; Vigneault-MacLean et al., 2007).

This study demonstrated that audio perception is influenced by both static and dynamic biases. For both azimuth and elevation judgements, the perceptual range was expanded compared to the sound sources’ veridical locations. Furthermore, contextual blocking resulted in additional bias, especially around the midline and horizon, compared to the full range condition. This illustrates that there is a warp in the mapping of real world sound sources and perceived location, and that the warp changes dynamically over time, as the brain recalibrates to represent the current environment.

## Acknowledgements

We would like to thank research assistants Sharon Wong and Prachetas Gumaste for help with data collection; software engineers Eli Stine, Christopher Poovey, and Alex Bezugly for development of the Unity app for running this experiment.

